# The potential of young vegetative quinoa (*Chenopodium quinoa*) as a new sustainable protein-rich winter leafy crop under Mediterranean climate

**DOI:** 10.1101/2023.08.01.551448

**Authors:** Lior Rubinovich, Reut Dagan, Yaron Lugasi, Shmuel Galili, Aviv Asher

## Abstract

The demand for protein products has significantly risen in the last few years. In western countries, animals are the primary source of protein; however, plants could take a share of this market due to lower production costs, among other advantages. Quinoa (*Chenopodium quinoa* Willd.) is a well-known but under-utilized protein-rich crop, commonly cultivated for grain production. These plants were recently evaluated for their use as a non-traditional, green leafy crop. Here we assessed the potential of young vegetative quinoa as a new sustainable winter leafy crop in Israel—serving as a model for Mediterranean semi-arid regions, by evaluating yield, protein content and quality. Five quinoa accessions were sown on three winter sowing dates over two consecutive years. Plants were harvested when they reached 10–13% dry matter (DM). DM yield ranged between 574 and 1,982 kg ha^-1^ and was generally higher in the second year. Protein content ranged from 14.4–34% and was generally higher in the first year. Protein yield ranged from 111–471 kg ha^-1^ and was greatest on the December sowing date. DM and protein yields were positively correlated with plant density. Protein content was negatively correlated with plant density and DM yield. Our findings show that 200 g DM of young vegetative quinoa can meet the protein and most essential amino acid requirements for a 70 kg human adult. Prospects for cultivating young vegetative quinoa in Mediterranean countries as a new sustainable, protein-rich winter leafy crop are therefore high, as supported by its high protein yields and quality, and its requirement for only scant irrigation. Further studies should examine economic and other agrotechnical parameters toward the geographical distribution and expansion of young vegetative quinoa cultivation.

## 1. Introduction

The demand for protein products has significantly risen in the last few years. The global protein ingredients market was valued at USD 38.5 billion in 2020 and is expected to expand at an annual growth rate of 10.5% from 2021 to 2028 [1]. A dietary shift in Western countries in the last century resulted in animals becoming the primary source of protein [2]. For instance, muscle meat protein content is relatively consistent across species, with an average near 22% and milk contains 3–7% protein, depending on the animal species [3]. However, plant proteins could take a share of the animal protein market (meat, egg and dairy) due to lower production costs, and their being perceived by consumers as being more ethical, healthy and environmentally friendly [4,5]. Grains, pulses, legumes, seeds and nuts are the main sources of plant proteins for human diets. The protein content (PC) of cereal grains ranges from 7 to 15% and is typically lower than that of animal protein sources on a dry matter (DM) basis [3]. Soybeans contain a PC of 35 to 40% and the PC of peas is generally about 25%, depending on the growing conditions and genotype. The PC of chickpeas, lentils, and beans is similar to peas, at around 20–36% [3]. The quinoa plant (*Chenopodium quinoa* Willd., Amaranthaceae) is a well-known but under-utilized protein-rich crop [6]. This pseudo-cereal originates from the South American Andes and is commonly grown for grain production for human consumption [7,8]. Quinoa is considered a climate-resilient crop and has a remarkable ability to grow in marginal environments [9,10]. Recently, the number of quinoa-producing countries has shown a rapid increase, and the crop is now grown commercially outside South America [11,12]. The quinoa grain’s PC is relatively high compared to the major cereal crops cultivated worldwide, such as wheat, corn and rice [13]; its high nutritional value is due to high levels of protein (11–19%) that contain all of the essential amino acids (EAA) [14,15]. It is also gluten-free, and may therefore be suitable for celiac patients [16,17]. Thus, quinoa is commonly considered a “superfood” or “functional food”, enhancing its economic attractiveness [18].

Many studies have shown the possibility of cultivating quinoa outside its area of origin in diverse agroecological regions, including semi-arid and arid climates [7,12]. Other studies have examined the effects of agrotechnical parameters such as row spacing, plant density, irrigation and fertilization on quinoa grain production [19–22]. Numerous recent studies have shown that quinoa can be used for purposes other than grain production. For example, several studies showed that the residual straw following grain harvest might be used as livestock feed, as it contains relatively high protein and mineral contents and is highly digestible [23–25]. Others showed that high quinoa hay biomass and nutritional quality suggest excellent prospects for its use as a high-quality ruminant feed [23,26,27]. Interestingly, the Incas and other earlier cultures included quinoa leaves in their diet to balance the lack of animal protein [28]. Indeed, in recent years, quinoa plants have also been evaluated for their use as a non-traditional, green leafy crop. The green leaves and tender stalks have high nutritional value and can be consumed raw or cooked, similar to, for instance, spinach [29,30]. Several studies have reported that the PC in quinoa leaves is greater than in quinoa grains, reaching over 25-37% in the DM, compared to 9.1-15.7% in grains [29,31,32]. The proximate composition of quinoa leaves’ crude fat, crude fiber, carbohydrates and energy is 2.4- 4.5%, 6.9-7.8%, 34% and 325 Kcal, respectively. In comparison, crude fat, crude fiber, carbohydrates and energy in the quinoa grains is 4-7.6%, 7-14.1%, 48.5-69.8% and 331-381 Kcal, respectively [29]. Quinoa leaves also contain higher PC than broccoli, spinach or chard (mangold) [31]. Moreover, quinoa protein contains all of the EAA, and the leaves have relatively low quantities of carbohydrates. Quinoa leaves also contain moderate levels of calcium, phosphorus, sodium and zinc, and high levels of copper, manganese and potassium [32]. Gawlik-Dziki et al. (2013) assessed the nutraceutical potential of quinoa leaves, and found that their extract contains significant amounts of various bioactive polyphenols linked with an inhibitory effect on prostate cancer cell proliferation [33]. Moreover, the content of saponins—bitter antinutritive compounds found in quinoa—is lower in the young leaves than in the grains [34,35]. The earliest detectable levels of sapogenins (a triterpenoid aglycone comprising about 50% of the saponin molecule) were discovered in quinoa leaves 82 days after sowing (DAS) [36]. Another study found the lowest saponin content in quinoa shoots at 60 DAS and the highest at 100 DAS [37].

Previous field trials have evaluated the yield of green leafy quinoa. A study conducted in Poland showed that quinoa should be harvested as a green leafy vegetable when it reaches a height of 20–30 cm. In spring and summer sowing, young quinoa’s average fresh weight biomass reached 317–1,389 and 245–777 g m^-2^, respectively [30]. In Egypt, young quinoa plants harvested 45 days after sowing had a greater fresh weight yield, 2.05–4.14 kg m^-2^ [38]. These findings indicate that growing quinoa as a green leafy vegetable has economic potential.

As there is only scarce data regarding the cultivation of young vegetative quinoa in Mediterranean countries, we aimed to investigate the yield and PC of this crop to evaluate its potential as a new protein-rich leafy crop in Israel. In particular, the objectives of this study were to examine the effects of different genotypes and sowing dates on young vegetative quinoa biomass yield, PC, protein yield and amino acid composition. We hypothesized that young quinoa’s aerial vegetative parts could be used as a new, sustainable, protein-rich leafy crop for Israel and other Mediterranean and semi-arid regions. Our findings show high prospects for cultivating young vegetative quinoa in Mediterranean countries as a new sustainable, protein-rich winter leafy crop.

## 2. Materials and Methods

### 2.1. Experimental site

Since Israel and other Mediterranean and semi-arid regions have an expensive and limited water supply, we investigated the potential of young vegetative quinoa cultivation as a winter crop, which is the rainy season in Israel. Plots were sown in November, December and January over two consecutive years to examine its potential cultivation under low amounts of irrigation. The experiments were conducted in northern Israel (The Avnei-Eitan research farm, altitude 375 masl, 32°81’64’’N 35°76’28’’E) from November 2020 to April 2022. The soil at the experimental site is of basaltic origin, composed of 60% clay, 35% silt and 5% sand with a pH of 7.6. The climate is Mediterranean, with a long-term annual average rainfall of 500–550 mm, mainly distributed between October and June. 92% of the total precipitation during the experimental season of 2020 to 2021 occurred between November 2020 and February 2021. Total precipitation during the growing season of 2021 to 2022 was slightly lower, with 95% of the total rainfall occurring from December 2021 to March 2022. Air temperature ranges were similar for the two seasons (Table 1).

**Table 1.**
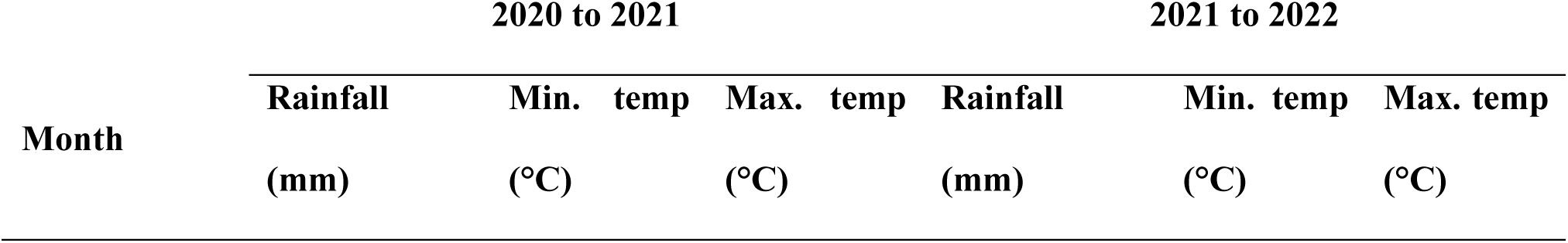
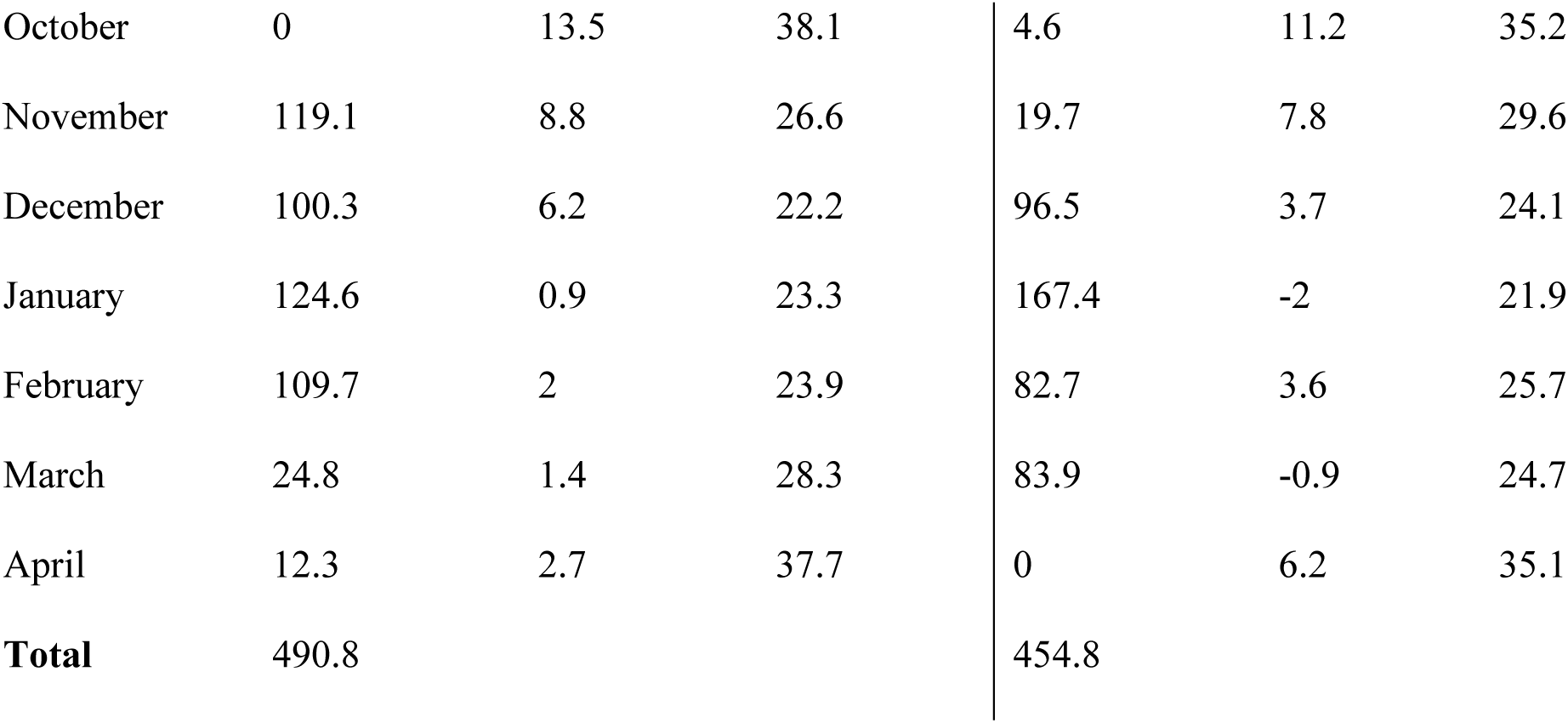
Monthly precipitation and temperatures at the Avnei-Eitan research farm during the 2020–2021 and 2021–2022 winter growing seasons (Israeli Meteorological Services).

### 2.2. Plant material and experimental design

Five quinoa accessions were used in the experiments: ‘Red Head’ (bright pink-red panicle with white seeds), ‘Mint Vanillà (white-panicle with white seeds), ‘Ivory’ (white-panicle with white seeds), ‘Oro de Vallè (golden-colored panicle with golden-brown seeds) and ‘Peppermint’ (white-panicle with white seeds; www.wildgardenseed.com, accessed 9.2023). These accessions were chosen for the experiment as they performed well in previous experiments conducted in the experimental region [20,23,39]. All accessions were obtained from Wild Garden Seed (Philomath, OR, USA). In all experiments, flowerbeds were prepared after cultivation and rolling. [20]Seeds were sown into six rows (16 cm between rows) at an intra-row density of 40–50 seeds m^-1^ using a manual seeder (a planned density of 240–300 seeds m^-2^). According to previous studies, this sowing density was chosen to achieve low stem diameter for enhanced palatability [20,30]. Quinoa accessions were sown into four 8-m^2^ (5 m long x 1.6 m wide) plots (repeats) in mid-November and mid-December 2020 and 2021, and mid-January 2021 and 2022, as different winter sowing dates may affect plant yield and quality parameters [23]. Plots were randomized on each sowing date. A single irrigation using sprinklers at a rate of 100 m^3^ ha^-1^ was applied only in plots that did not receive rain within 10 days after sowing. The plots were fertilized by 92 kg N ha^-1^ as urea (46% N). All plots were manually weeded once during cultivation.

### 2.3. Plant density and yield measurements

To determine plant density at harvest (plants m^−2^), plants were counted in a representative 1-m^2^ area in each plot. The representative area in each plot was determined by an external evaluation of the plant density and appearance of the plants. From 50 DAS, five plants with similar external characteristics were collected every three days and weighed before and after drying at 60 °C for 48 h and the %DM was calculated. When these plants reached 10% DM, the aerial parts of all the plants in each plot were hand-harvested ∼5 cm above the ground and weighed for fresh matter yield. Harvest timing was set to avoid saponin accumulation and according to previous experiments [36,38,40]. To determine plant %DM, representative samples of each plot were collected and weighed before and after drying at 60 °C for 48 h. Plot DM yield was calculated as fresh matter yield x plant %DM.

### 2.4. Plant protein content, protein yield and amino acid composition

Using the Kjeldahl method [32], each quinoa sample’s PC (% of DM) was determined at the Milouda and Migal laboratories in Kiryat Shmona, Israel. Plot protein yield was calculated as DM yield x PC. The amino acid composition was determined at the Merieux NutriScience laboratories in Resana, Italy. Briefly, the samples were subjected to hot enzymatic hydrolysis of the proteins. The solution obtained from hydrolysis was diluted with deionized water and methanol and analyzed by HPLC.

### 2.5. Statistical analysis

The data are presented as mean value ± standard error (SE). The effects of the different accessions on plant density, DM yield, PC and protein yield were analyzed separately for each year’s sowing date. The effects of the different sowing dates on the amino acid composition were analyzed separately for each amino acid each year. Results were subjected to one-way analysis of variance (ANOVA) followed by Tukey’s HSD test (to compare all pairs) using JMP software version 11.0.0 (SAS Institute, Cary, NC, USA). The significance between treatments was checked at P < 0.05. Results of plant density, DM yield, PC and protein yield were also subjected to a two-tailed Pearson correlation matrix using Prism software version 9.0.0 (GraphPad Software, San Diego, CA, USA). Correlations were calculated and regression lines were drawn for DM yield, PC, protein yield and Plant density, as well as PC and DM yield. Data points from all plots during the two years of the experiments were subjected to the correlation analysis.

## 3. Results

### 3.1. Plant density

In the first year, plants from the plots sown in November 2020, December 2020 and January 2021 were harvested 70, 73 and 69 days after sowing, respectively (Fig 1).

**Fig. 1.**
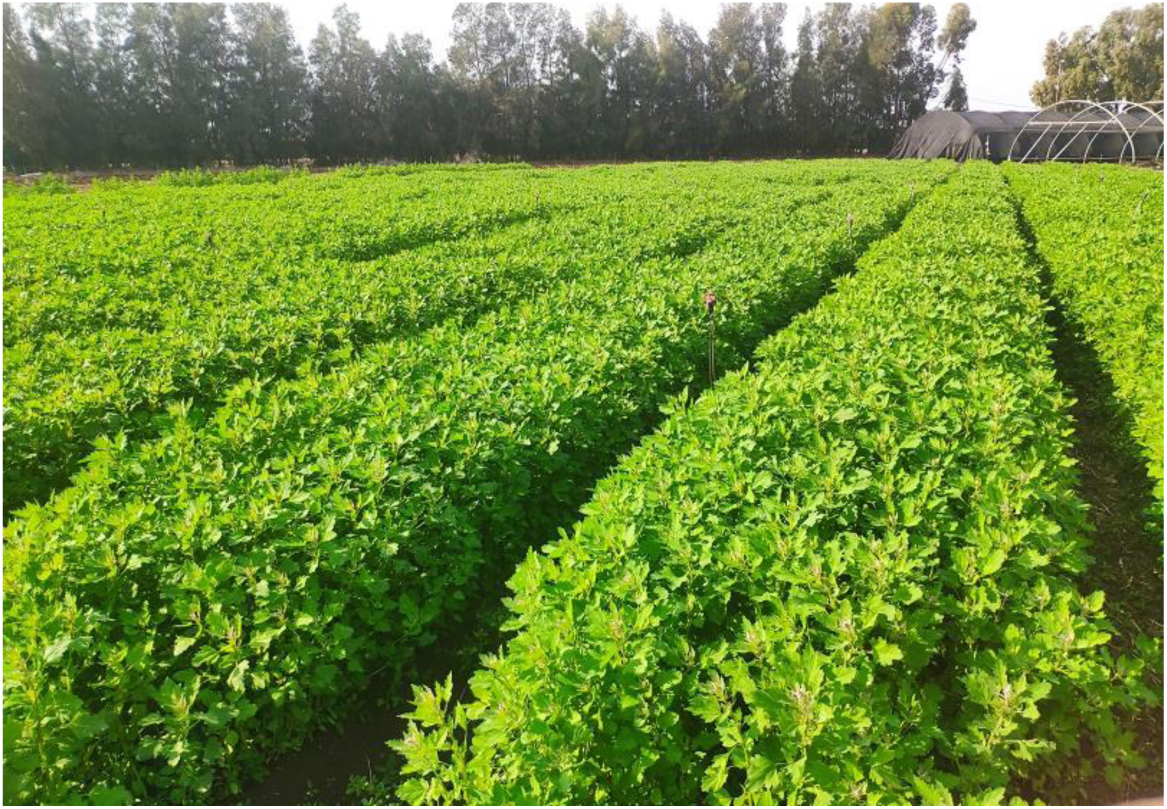
Young vegetative quinoa. A representative photograph of young vegetative quinoa grown at the Avnei-Eitan research farm before harvest.

Actual plant densities at harvest are shown in Fig 2. In plots sown in November 2020, plant density ranged between 90 (accession Oro de Valle) and 192 (‘Peppermint’) plants m^-2^. In plots sown in December 2020, plant density ranged between 179 and 264 plants m^-2^ (accessions Mint Vanilla and Red Head, respectively). There were no differences (*P* > 0.05) between plant densities of the different accessions from these sowing dates. In plots sown in January 2021, plant density ranged between 163 and 302 plants m^-2^. Accession Red Head had the highest plant density, significantly (*P* < 0.05) higher than ‘Oro de Vallè. In the second year, plants from the plots sown in November 2021, December 2021 and January 2022 were harvested 99, 107 and 88 days after sowing, respectively (Fig 2a). In plots sown in November 2021, plant density ranged between 150 and 333 plants m^-2^. Accession Oro de Valle had the highest plant density, significantly (*P* < 0.05) higher than ‘Mint Vanillà and ‘Ivory’. In plots sown in December 2021, plant density ranged between 330 and 521 plants m^-2^. Accession Mint Vanilla had the highest plant density, significantly (*P* < 0.05) higher than ‘Red Head’, ‘Oro de Vallè and ‘Peppermint’. In plots sown in January 2022, plant density ranged between 142 and 567 plants m^-2^. Accession Mint Vanilla had the highest plant density, significantly (*P* < 0.05) higher than all other accessions (Fig 2b).

**Fig. 2.**
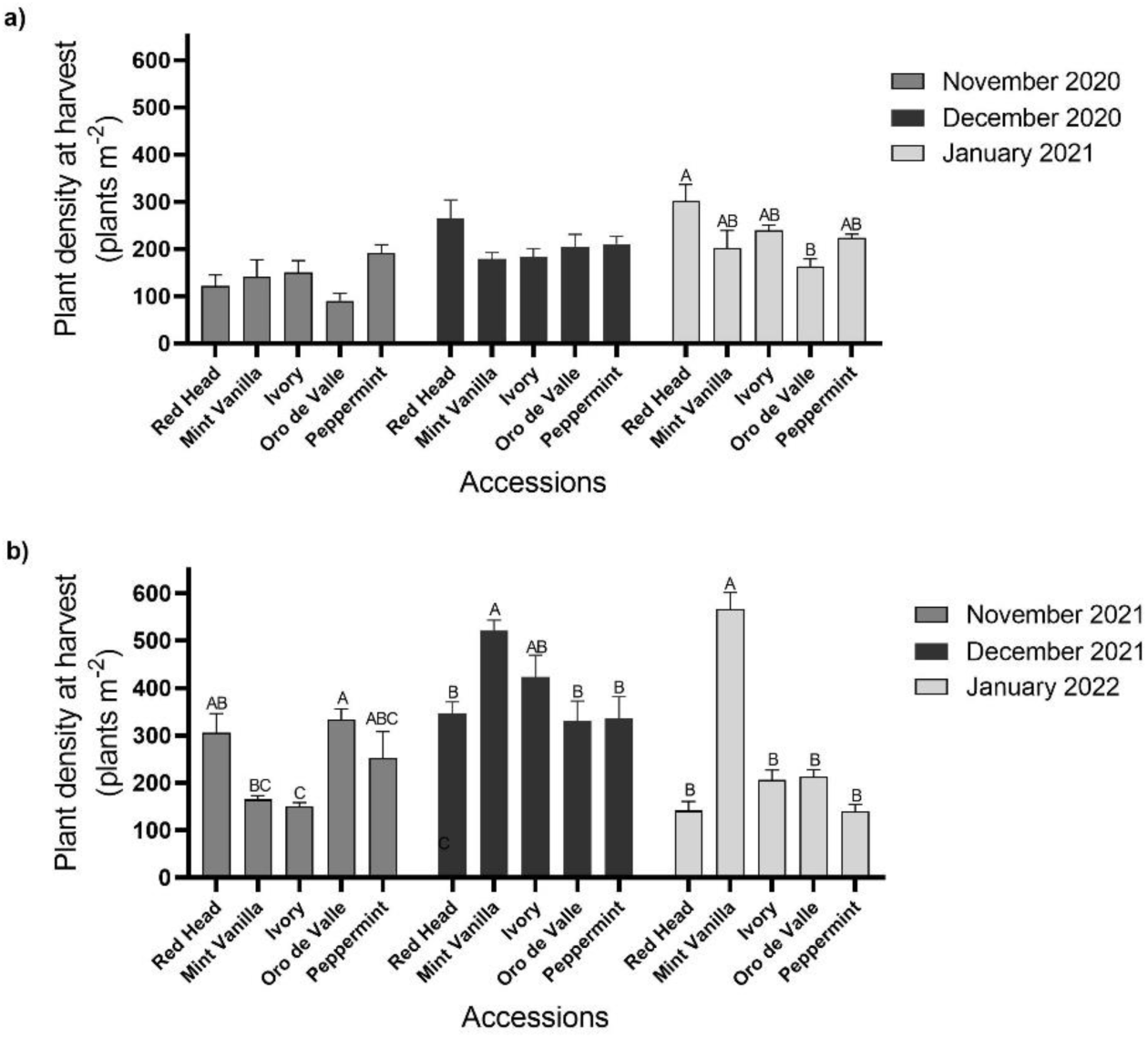
Young vegetative quinoa plant density. Quinoa was sown in the Avnei-Eitan research farm during the winters of 2020–2021 and 2021–2022. Results show plant density at harvest in plots sown in November 2020, December 2020, and January 2021 (a); and in November 2021, December 2021, and January 2022 (b). Results are presented as mean value ± standard error of four 5-m^2^ sample plots per accession on each sowing date; columns marked with different letters differ significantly by Tukey-HSD (*P* < 0.05).

### 3.2. Dry matter yield

Quinoa DM yield is shown in Fig 3. In plots sown in November 2020, DM yield of young vegetative quinoa ranged between 572 and 1200 kg DM ha^-1^ (accessions Oro de Valle and Peppermint, respectively). In plots sown in December 2020, DM yield ranged between 693 and 1069 kg DM ha^-1^ (accessions Oro de Valle and Mint Vanilla), respectively. There were no differences (*P* > 0.05) between the DM yields of the different accessions on November and December 2020 sowing dates. In plots sown in January 2021, DM yield ranged between 701 and 1174 kg DM ha^-1^. Accession Peppermint had the highest DM yield, significantly higher (*P* < 0.05) than ‘Oro de Valle’ (Fig 3a). In the second year (Fig 3b), in plots sown in November and December 2021 and January 2022, DM yield ranged between 1088 and 1434 kg DM ha^-1^ (‘Red Head’ and ‘Oro de Valle’), 1477 and 1982 kg DM ha^-1^ (‘Red Head’ and ‘Ivory’), and 618 and 1384 kg DM ha^-1^ (‘Red Head’ and ‘Ivory’), respectively. There were no significant differences (*P* > 0.05) between the DM yields of the different accessions for any of these sowing dates.

**Fig. 3.**
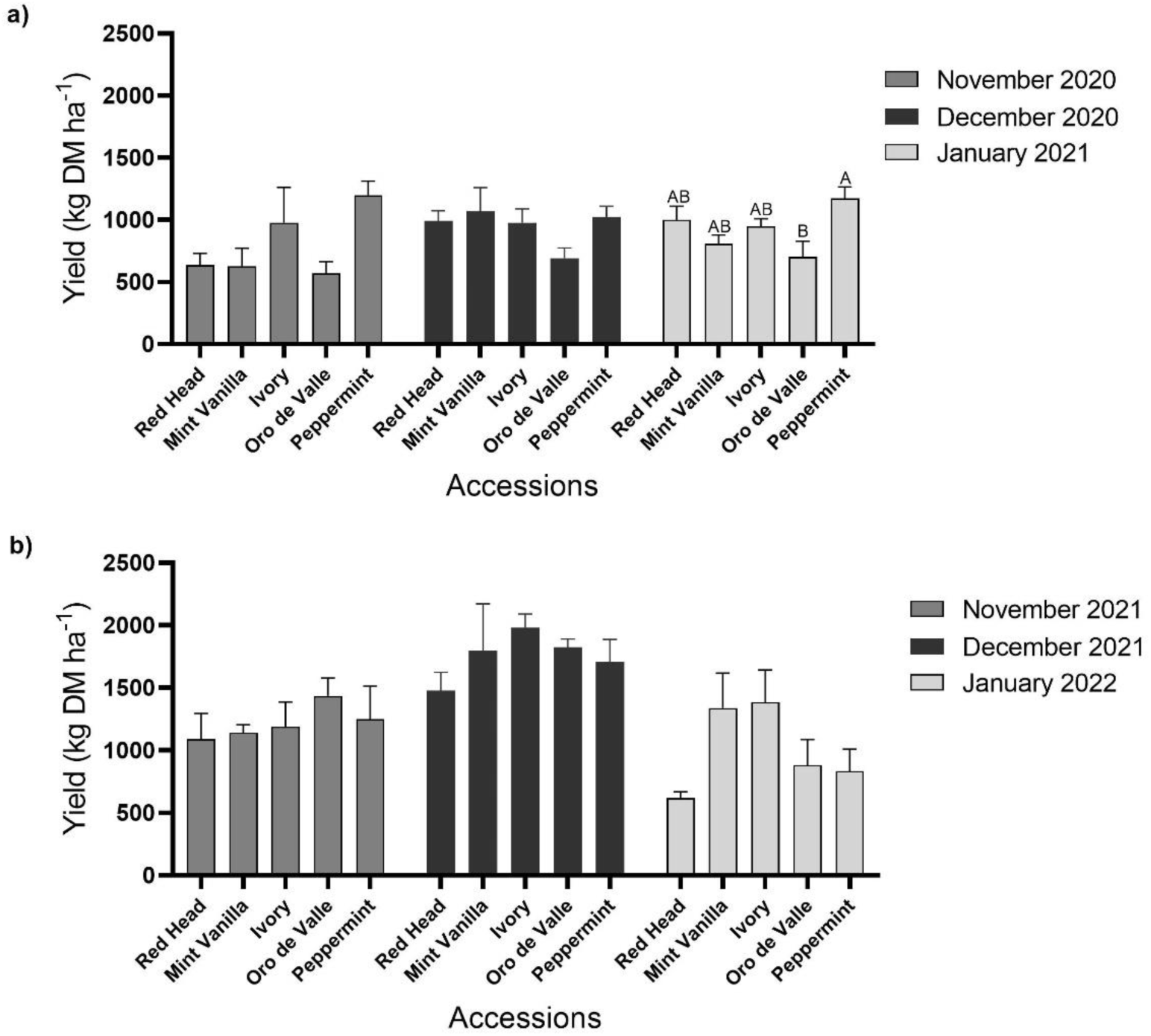
Young vegetative quinoa yield. Quinoa was sown in the Avnei-Eitan research farm during the winters of 2020–2021 and 2021–2022. Results show DM yield in plots sown in November 2020, December 2020 and January 2021 (a); and in November 2021, December 2021, and January 2022 (b). Results are presented as mean value ± standard error of four 5-m^2^ sample plots per accession on each sowing date; columns marked with different letters differ significantly by Tukey-HSD (*P* < 0.05).

### 3.3. Protein content and protein yield

Quinoa PC is shown in Fig 4. In plots sown in November 2020, the PC of young vegetative quinoa ranged between 30.6 and 32.4% (accessions Peppermint and Red Head, respectively). In plots sown in December 2020, quinoa PC ranged between 28.2 and 34% (‘Red Head’ and ‘Mint Vanilla’, respectively). In plots sown in January 2021, quinoa PC ranged between 24.8 and 29.2% (‘Ivory’ and ‘Mint Vanilla’, respectively) (Fig 4a). In the second year (Fig 4b), in plots sown in November 2021, quinoa PC ranged between 25.9 and 28.3% (‘Oro de Valle’ and ‘Red Head’, respectively). In plots sown in December 2021, quinoa PC ranged between 19.4 and 23.7% (‘Mint Vanilla’ and ‘Ivory’, respectively). In plots sown in January 2022, quinoa PC ranged between 14.4 and 19.4% (‘Ivory’ and ‘Peppermint’, respectively). There were no significant differences (*P* > 0.05) between the PC of the different accessions for any of these sowing dates.

**Fig. 4.**
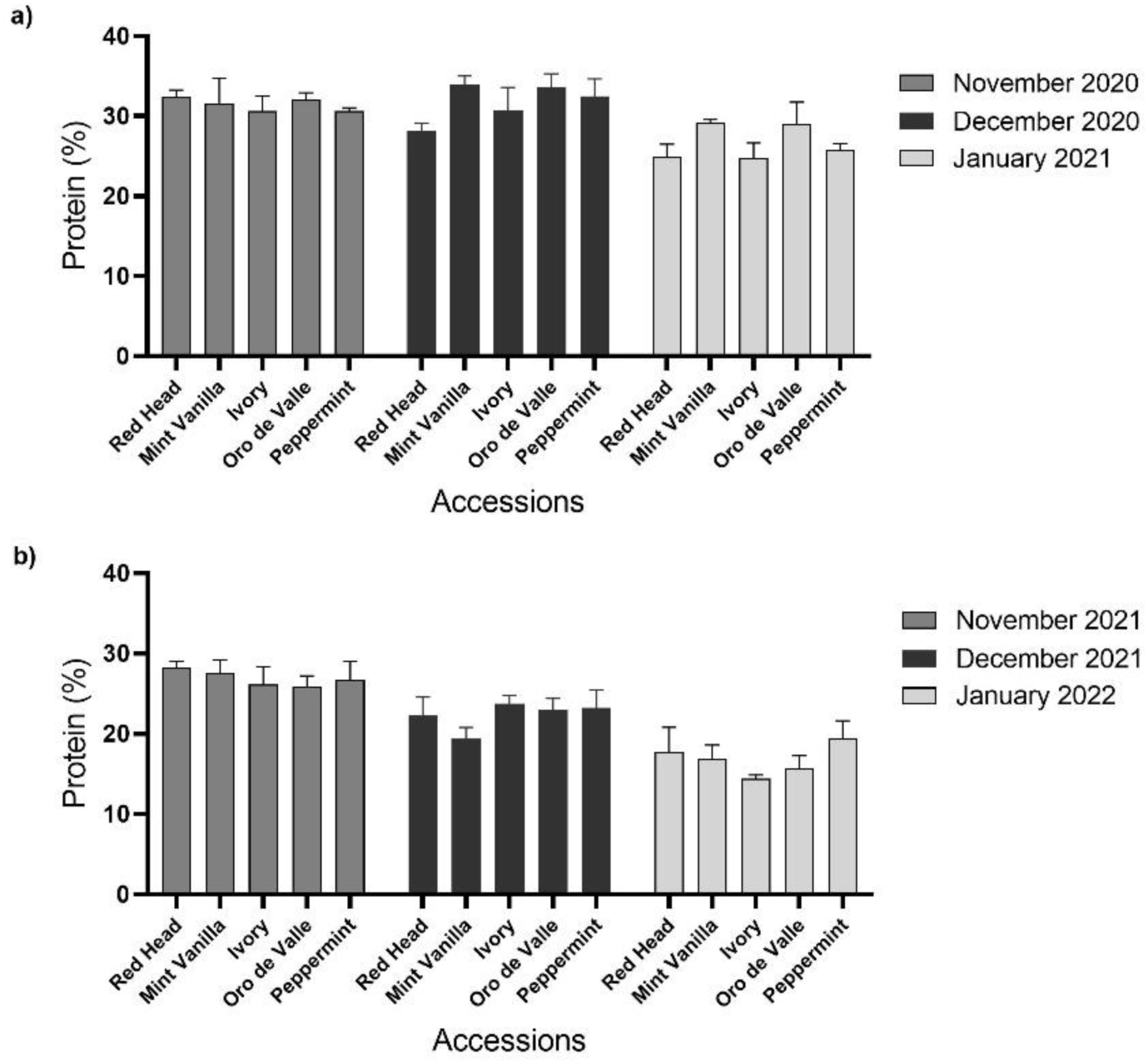
Young vegetative quinoa protein content. Quinoa was sown in the Avnei-Eitan research farm during the winters of 2020–2021 and 2021–2022. Results show protein content (% DM) in quinoa plants sown in November 2020, December 2020, and January 2021 (a); and in November 2021, December 2021, and January 2022 (b). Results are presented as mean value ± standard error of four 5-m^2^ sample plots per accession on each sowing date.

Quinoa protein yield is shown in Fig 5. In plots sown in November 2020, protein yield ranged between 182 and 366 kg ha^-1^ (accessions Oro de Valle and Peppermint, respectively). In plots sown in December 2020, protein yield ranged between 230 and 359 kg ha^-1^ (‘Oro de Valle’ and ‘Mint Vanilla’, respectively). In plots sown in January 2021, protein yield ranged between 201 and 303 kg ha^-1^ (‘Oro de Valle’ and ‘Peppermint’, respectively) (Fig 5a). In the second year (Fig 5b), in plots sown in November 2021, protein yield ranged between 303 and 367 kg ha^-1^ (‘Ivory’ and ‘Oro de Valle’, respectively). In plots sown in December 2021, protein yield ranged between 330 and 471 kg ha^-1^ (‘Red Head’ and ‘Ivory’, respectively). In plots sown in January 2022, protein yield ranged between 111 and 240 kg ha^-1^ (‘Red Head’ and ‘Mint Vanilla’, respectively). There were no significant differences (*P* > 0.05) between the protein yields of the different accessions for each of these sowing dates.

**Fig. 5.**
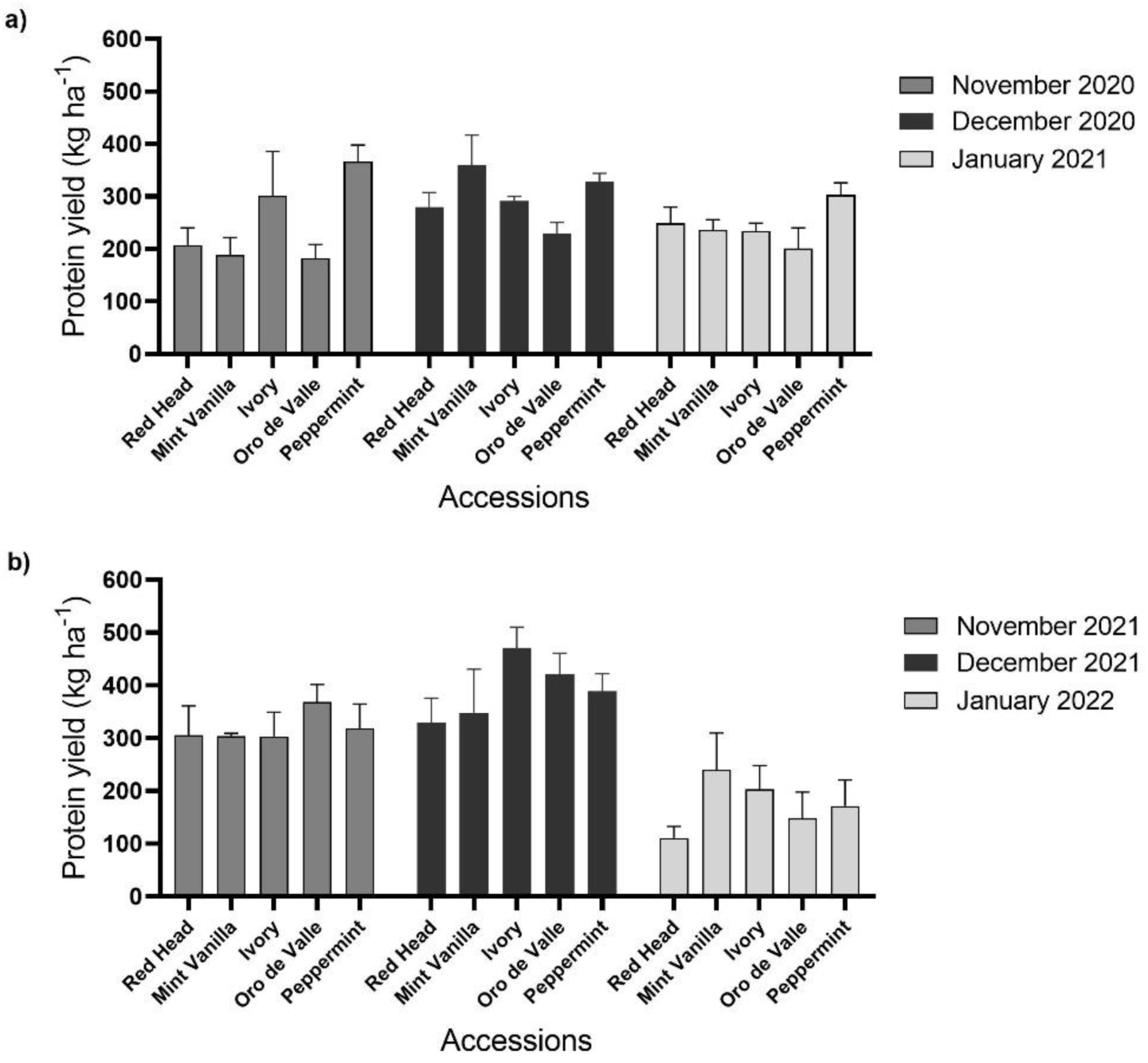
Young vegetative quinoa protein yield. Quinoa was sown in the Avnei-Eitan research farm during the winters of 2020–2021 and 2021–2022. Results show protein yield in quinoa plants sown in November 2020, December 2020 and January 2021 (a); and in November 2021, December 2021, and January 2022 (b). Results are presented as mean value ± standard error of four 5-m^2^ sample plots per accession on each sowing date.

### 3.4. Correlation between traits

The correlations between DM yield, PC and protein yield with plant density and the correlation between PC and DM yield of the young vegetative quinoa are shown in Fig 6. They are based on data from both experimental years. Young vegetative quinoa DM yield was positively correlated (r = 0.616, *P* < 0.0001) with plant density (Fig 6a), but an opposite and low correlation was found between this parameter and PC (r = -0.376, *P* < 0.0001) (Fig 6b). A positive moderate correlation (r = 0.43, *P* < 0.0001) was found between protein yield and plant density (Fig. 6c), which was opposite to the low negative correlation between PC and plant density. The correlation between PC and DM yield was negative (r = -0.311, *P* < 0.001) (Fig 6d), similar to the negative correlation between PC and plant density.

**Fig. 6.**
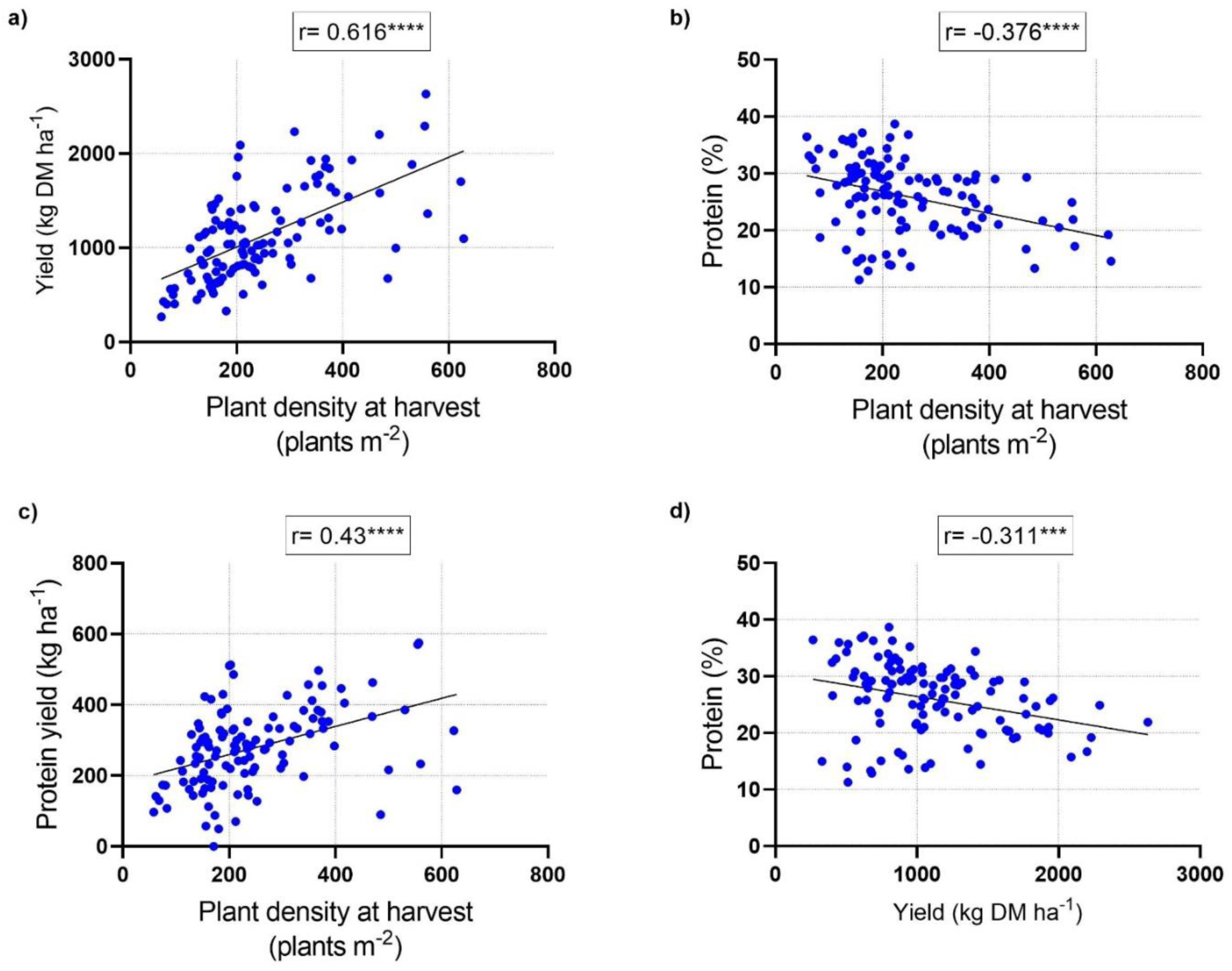
Correlations of young vegetative quinoa yield parameters. Pearson’s correlation coefficients between yield and plant density at harvest (a), protein content and plant density at harvest (b), protein yield and plant density at harvest (c), and protein content and yield (d). Values represent measurements from all plots from the winters of 2020–2021 and 2021–2022; each value represents measurements from one plot. The correlation coefficient value (r) is shown for each graph along with the significance of the Pearson correlation values (\*\*\**P* < 0.001, \*\*\*\**P* < 0.0001).

### 3.5. Amino acid composition

As accession Peppermint had relatively high protein yield values during the first year of the experiment, it was chosen for amino acid composition analysis. During the first year of the investigation, the young vegetative quinoa contained all of the EAA (Table 2). Among them, the quinoa was characterized by a high content (g per 100 g DM) of leucine and lysine. Their levels were significantly (*P* < 0.05) higher in plots sown in December 2020 compared to November 2020. Tryptophan levels were significantly (*P* < 0.05) higher in plots sown in December 2020 and January 2021 compared to November 2020. There were no significant differences (*P* > 0.05) in the other EAA levels between the different sowing dates. For comparison, the recommended daily EAA intake for humans (g per 70 kg body weight) is shown in Table 2. On the different sowing dates, histidine, isoleucine, leucine, lysine, methionine, phenylalanine (with tyrosine), threonine, tryptophan and valine levels (g per 100 g DM) in the young vegetative quinoa ranged from 50–63%, 41–57%, 53–68%, 60–112%, 37–50%, 79–110%, 84–119%, 70–164% and 53–63% of the recommended daily intake, respectively. Among the non-essential amino acids (NEAA), young vegetative quinoa accession Peppermint was characterized by high glutamic and aspartic acid contents. Aspartic acid and serine levels were significantly (*P* < 0.05) higher in plots sown in December 2020 and January 2021 compared to November 2020. Glutamic acid and glycine levels were significantly (*P* < 0.05) higher in plots sown in December 2020 compared to November 2020 and January 2021. Alanine levels were significantly (*P* < 0.05) higher in plots sown in December 2020 compared to November 2020. In the experiment’s first year, there were no significant differences (*P* > 0.05) in the other NEAA levels between the different sowing dates (Table 2).

**Table 2.**
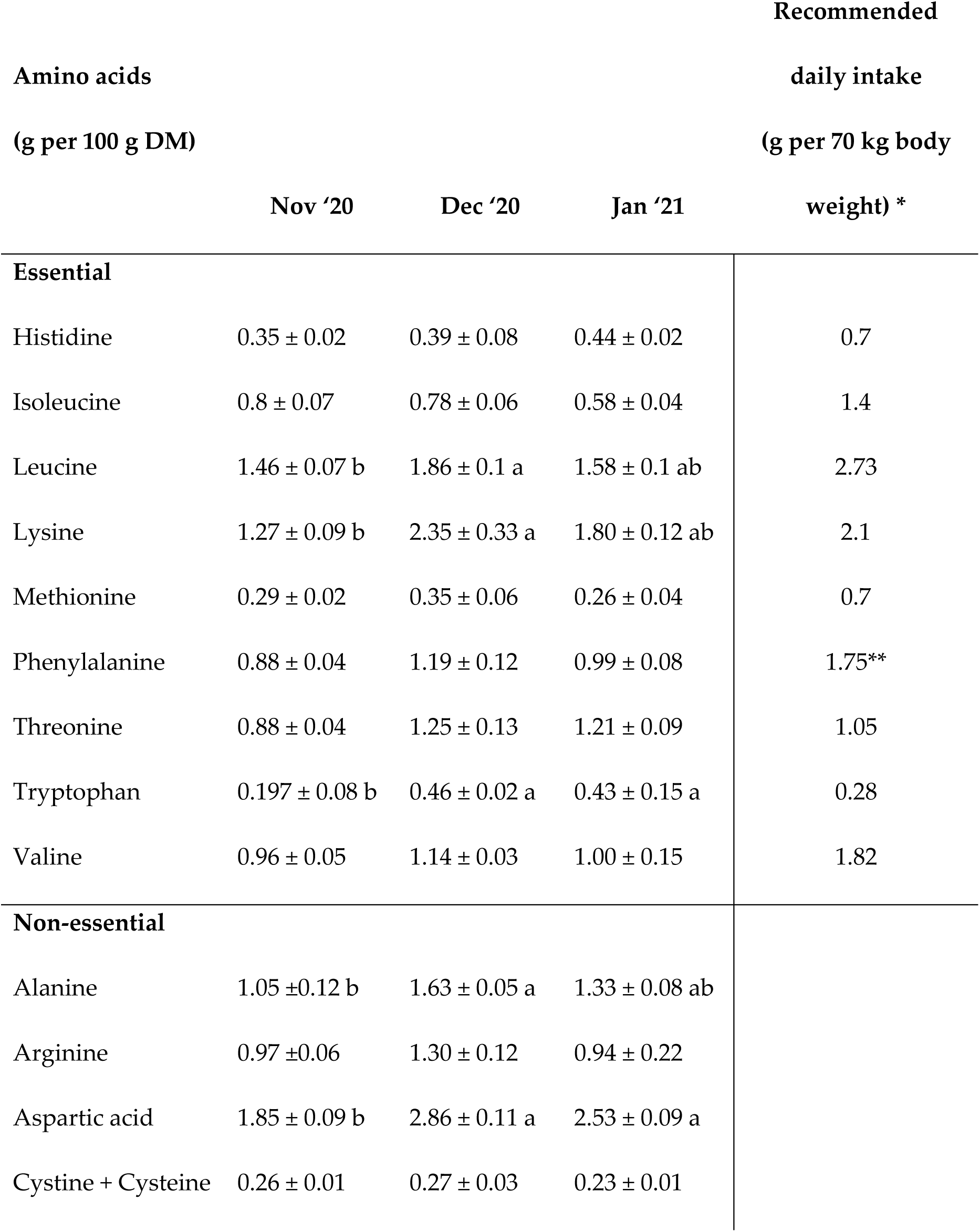
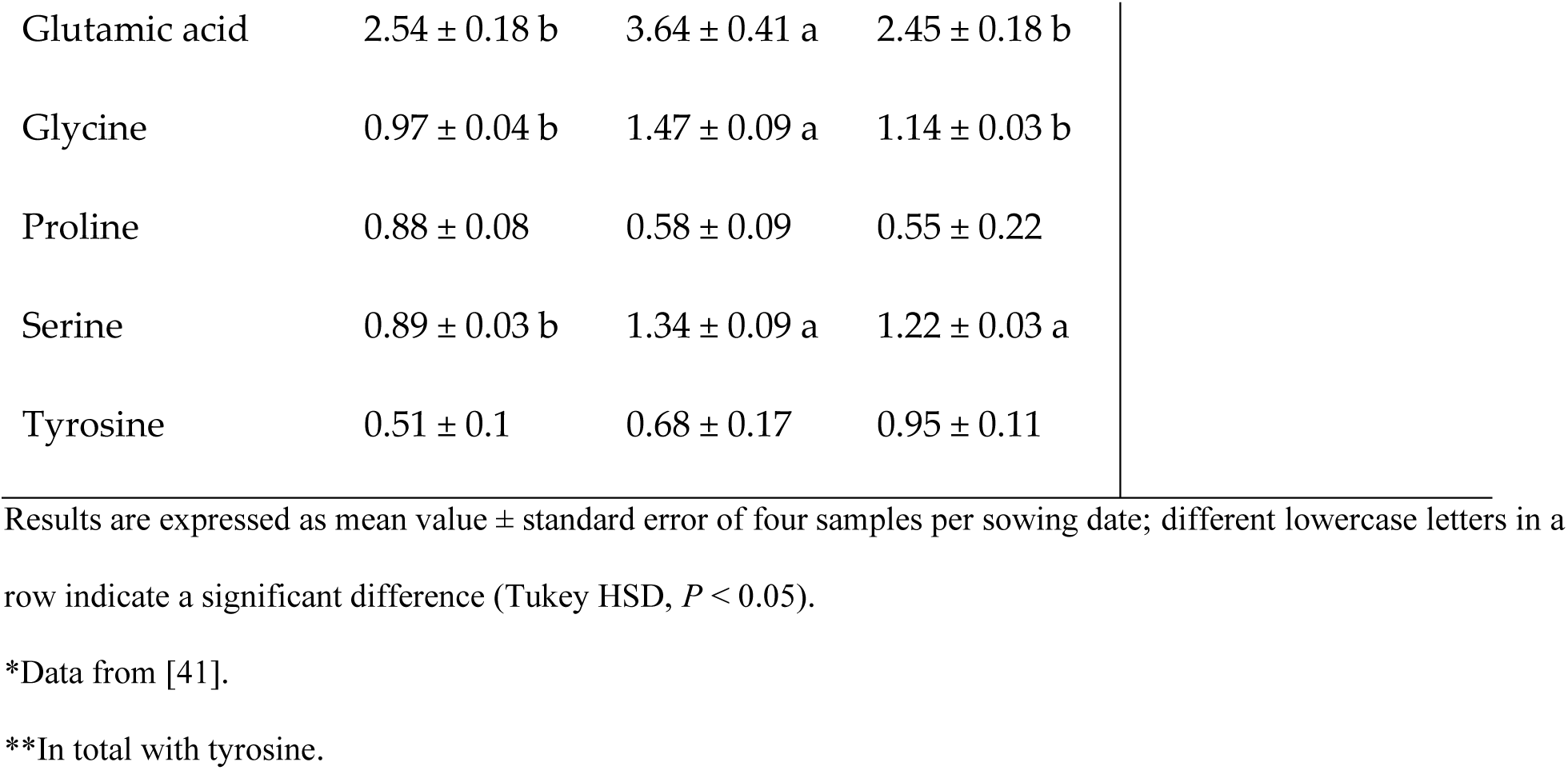
Amino acid composition (g per100 g DM) of accession Peppermint from the 2020–2021 winter sowing dates: November 2020, December 2020, and January 2021.

| Amino acids<br>(g per 100 g DM) |  |  |  | Recommended<br>daily intake<br>(g per 70 kg body<br>weight) * |
| --- | --- | --- | --- | --- |
|  | Nov '20 | Dec '20 | Jan '21 |  |
| **Essential** |  |  |  |  |
| Histidine | 0.35 ± 0.02 | 0.39 ± 0.08 | 0.44 ± 0.02 | 0.7 |
| Isoleucine | 0.8 ± 0.07 | 0.78 ± 0.06 | 0.58 ± 0.04 | 1.4 |
| Leucine | 1.46 ± 0.07 b | 1.86 ± 0.1 a | 1.58 ± 0.1 ab | 2.73 |
| Lysine | 1.27 ± 0.09 b | 2.35 ± 0.33 a | 1.80 ± 0.12 ab | 2.1 |
| Methionine | 0.29 ± 0.02 | 0.35 ± 0.06 | 0.26 ± 0.04 | 0.7 |
| Phenylalanine | 0.88 ± 0.04 | 1.19 ± 0.12 | 0.99 ± 0.08 | 1.75** |
| Threonine | 0.88 ± 0.04 | 1.25 ± 0.13 | 1.21 ± 0.09 | 1.05 |
| Tryptophan | 0.197 ± 0.08 b | 0.46 ± 0.02 a | 0.43 ± 0.15 a | 0.28 |
| Valine | 0.96 ± 0.05 | 1.14 ± 0.03 | 1.00 ± 0.15 | 1.82 |
| **Non-essential** |  |  |  |  |
| Alanine | 1.05 ± 0.12 b | 1.63 ± 0.05 a | 1.33 ± 0.08 ab |  |
| Arginine | 0.97 ± 0.06 | 1.30 ± 0.12 | 0.94 ± 0.22 |  |
| Aspartic acid | 1.85 ± 0.09 b | 2.86 ± 0.11 a | 2.53 ± 0.09 a |  |
| Cystine + Cysteine | 0.26 ± 0.01 | 0.27 ± 0.03 | 0.23 ± 0.01 |  |

|  |  |  |  |
| --- | --- | --- | --- |
| Glutamic acid | 2.54 ± 0.18 b | 3.64 ± 0.41 a | 2.45 ± 0.18 b |
| Glycine | 0.97 ± 0.04 b | 1.47 ± 0.09 a | 1.14 ± 0.03 b |
| Proline | 0.88 ± 0.08 | 0.58 ± 0.09 | 0.55 ± 0.22 |
| Serine | 0.89 ± 0.03 b | 1.34 ± 0.09 a | 1.22 ± 0.03 a |
| Tyrosine | 0.51 ± 0.1 | 0.68 ± 0.17 | 0.95 ± 0.11 |

Results are expressed as mean value ± standard error of four samples per sowing date; different lowercase letters in a row indicate a significant difference (Tukey HSD, $P < 0.05$ ).
\*Data from [41].
\*\*In total with tyrosine.

During the second year of the experiment, the young vegetative quinoa also contained all EAA and was characterized by a high content (g per 100 g DM) of leucine and lysine (Table 3). However, there were no significant differences (*P* > 0.05) in EAA levels between the different sowing dates. On the different sowing dates, histidine, isoleucine, leucine, lysine, methionine, phenylalanine (with tyrosine), threonine, tryptophan and valine levels (g per 100 g DM) in the young vegetative quinoa, ranged from 53–66%, 49–72%, 52–65%, 50–60%, 46–54%, 87–103%, 77–98%, 54–66% and 52–66% of the recommended daily EAA intake (Table 3), respectively. Among the NEAA, young vegetative quinoa was characterized by relatively high glutamic acid, aspartic acid and arginine contents. Alanine, arginine and serine levels were significantly (*P* < 0.05) higher in plots sown in November 2021 compared to January 2022. Glutamic acid levels were significantly (*P* < 0.05) higher in plots sown in November 2021 compared to December 2021 and January 2022. There were no significant differences (*P* > 0.05) in the other NEAA levels between the different sowing dates of the second year of the experiment (Table 3).

**Table 3.**
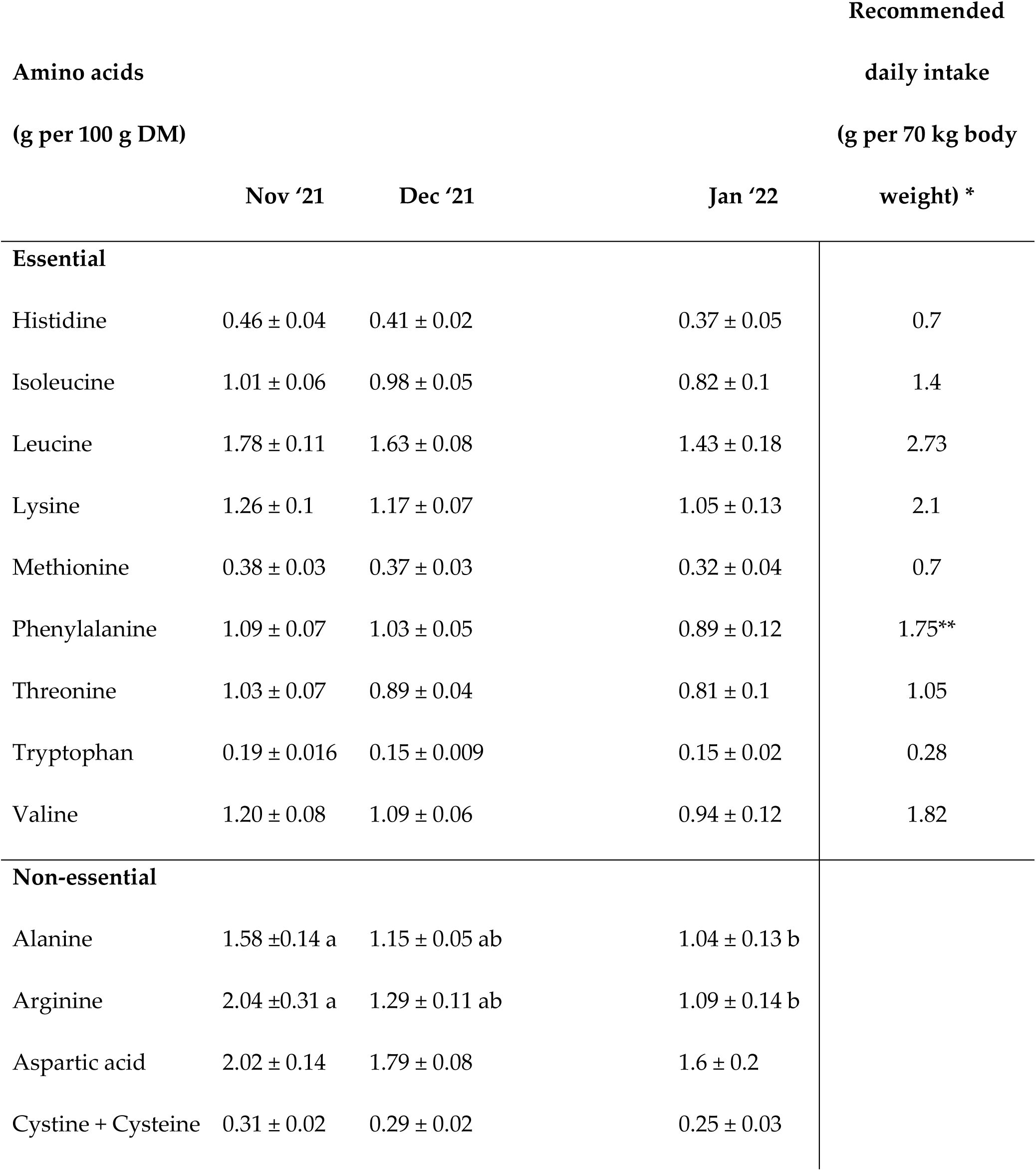
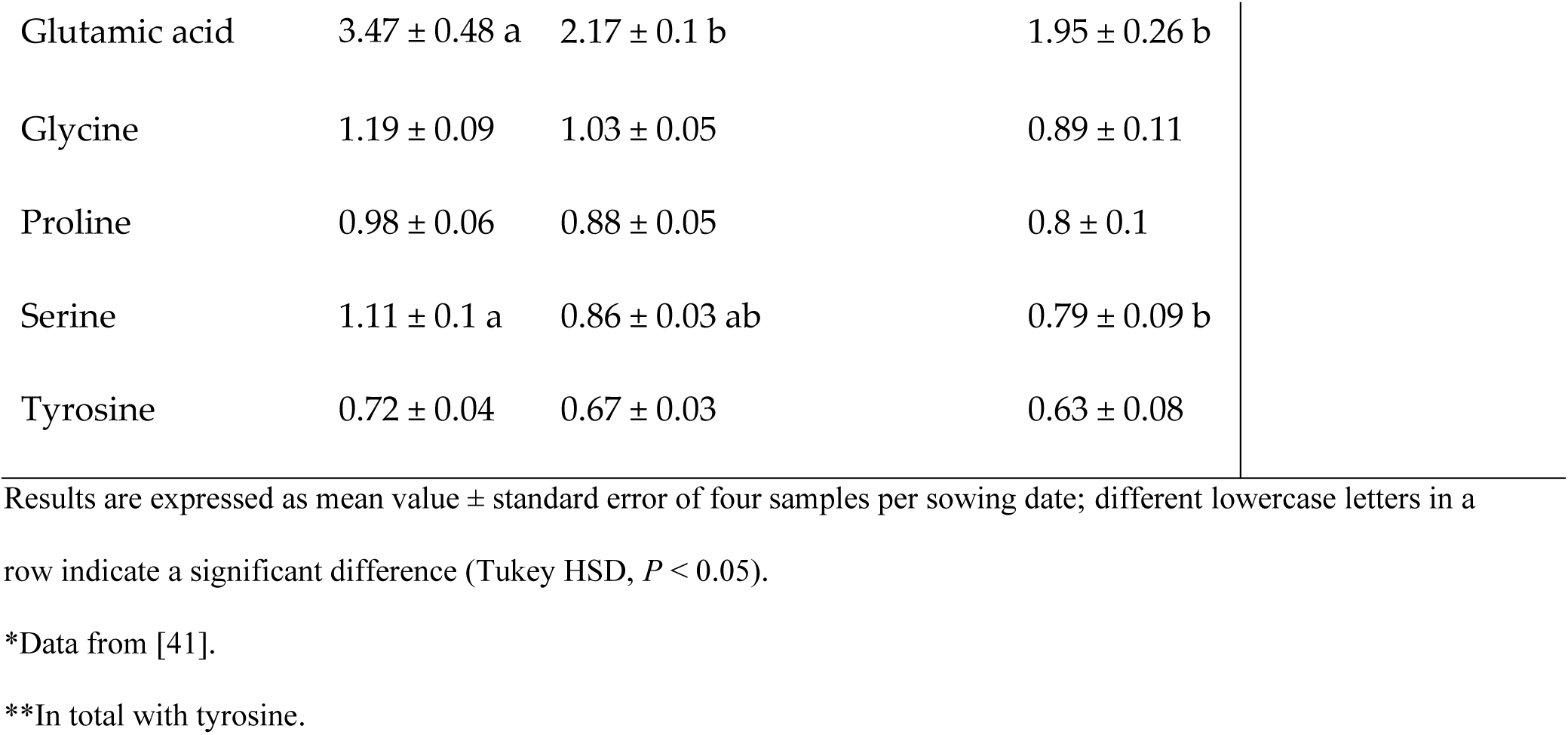
Amino acid composition (g per100 g DM) of accession Peppermint from the 2021–2022 winter sowing dates: November 2021, December 2021, and January 2022.

| Amino acids<br>(g per 100 g DM) |  |  |  | Recommended<br>daily intake<br>(g per 70 kg body<br>weight) * |
| --- | --- | --- | --- | --- |
|  | Nov '21 | Dec '21 | Jan '22 |  |
| **Essential** |  |  |  |  |
| Histidine | 0.46 ± 0.04 | 0.41 ± 0.02 | 0.37 ± 0.05 | 0.7 |
| Isoleucine | 1.01 ± 0.06 | 0.98 ± 0.05 | 0.82 ± 0.1 | 1.4 |
| Leucine | 1.78 ± 0.11 | 1.63 ± 0.08 | 1.43 ± 0.18 | 2.73 |
| Lysine | 1.26 ± 0.1 | 1.17 ± 0.07 | 1.05 ± 0.13 | 2.1 |
| Methionine | 0.38 ± 0.03 | 0.37 ± 0.03 | 0.32 ± 0.04 | 0.7 |
| Phenylalanine | 1.09 ± 0.07 | 1.03 ± 0.05 | 0.89 ± 0.12 | 1.75** |
| Threonine | 1.03 ± 0.07 | 0.89 ± 0.04 | 0.81 ± 0.1 | 1.05 |
| Tryptophan | 0.19 ± 0.016 | 0.15 ± 0.009 | 0.15 ± 0.02 | 0.28 |
| Valine | 1.20 ± 0.08 | 1.09 ± 0.06 | 0.94 ± 0.12 | 1.82 |
| **Non-essential** |  |  |  |  |
| Alanine | 1.58 ± 0.14 a | 1.15 ± 0.05 ab | 1.04 ± 0.13 b |  |
| Arginine | 2.04 ± 0.31 a | 1.29 ± 0.11 ab | 1.09 ± 0.14 b |  |
| Aspartic acid | 2.02 ± 0.14 | 1.79 ± 0.08 | 1.6 ± 0.2 |  |
| Cystine + Cysteine | 0.31 ± 0.02 | 0.29 ± 0.02 | 0.25 ± 0.03 |  |

|  |  |  |  |
| --- | --- | --- | --- |
| Glutamic acid | 3.47 ± 0.48 a | 2.17 ± 0.1 b | 1.95 ± 0.26 b |
| Glycine | 1.19 ± 0.09 | 1.03 ± 0.05 | 0.89 ± 0.11 |
| Proline | 0.98 ± 0.06 | 0.88 ± 0.05 | 0.8 ± 0.1 |
| Serine | 1.11 ± 0.1 a | 0.86 ± 0.03 ab | 0.79 ± 0.09 b |
| Tyrosine | 0.72 ± 0.04 | 0.67 ± 0.03 | 0.63 ± 0.08 |

Results are expressed as mean value ± standard error of four samples per sowing date; different lowercase letters in a row indicate a significant difference (Tukey HSD, $P < 0.05$ ).
\*Data from [41].
\*\*In total with tyrosine.

## 4. Discussion

We assessed the potential of young vegetative quinoa as a new winter leafy crop in Israel and as a model for Mediterranean semi-arid regions by evaluating yield, PC and quality. Five quinoa accessions were sown on three winter sowing dates over two consecutive years. The greatest yield (1,982 kg DM ha^-1^) was achieved with accession Ivory sown in December 2021 (Fig 3). The cultivation of quinoa as a leafy crop is still rare worldwide. Therefore, there are no comparable data on its production parameters from the Food and Agriculture Organization of the United Nations (FAO). However, the fresh yield of quinoa (∼17,300 kg ha^-1^ with accession Ivory sown in December 2021) was relatively high compared to the commercial yields of spinach, a more common leafy crop. According to the FAO, the fresh yield of commercial spinach in Italy and Egypt from 2,018 to 2,021 ranged between 15,558 and 16,574 kg ha^-1^ and 14,952 and 16,411 kg ha^-1^, respectively (www.fao.org, accessed 2.2023). However, it is essential to note that because crops are given more care and attention in small-scale field studies, yields are often higher than in commercial-scale production. Even so, the quinoa yield obtained in this study was higher than that reported in earlier small-scale field experiments. For example, a recent study showed that under saline and non-saline conditions in Egypt, quinoa fresh vegetative yield reached 4,470 and 5,985 kg ha^-1^, respectively [40]. In another study conducted in India, the fresh yield of the first harvest of quinoa foliage reached between 6,240 and 4,100 kg ha^-1^ in two consecutive seasons [42]. Interestingly, in that same study, the reported cumulative fresh yield of three consecutive harvests of the same plots reached 18,999 and 13,900 kg ha^-1^ in the two respective seasons of the experiment. Thus, quinoa yield may be significantly increased by growing the plants after the first harvest and harvesting them at intervals, similar to spinach, alfalfa, elephant grass, and other leafy field crops [43–45]. However, multiple harvests may eventually reduce quinoa plant yield and quality [42]. Another suggested possibility for maximizing quinoa yield while retaining quality is to establish a new cultivation on the same plot following harvest, similar to spinach and rocket [45]. However, as pests and diseases may develop with high plant density and re-seeding of the same crop on the same plot, crop rotation should be considered to reduce possible adverse effects [46].

In this study, PC in young vegetative quinoa ranged between 14.4% and 34%, and in general, it was higher in the first year of the experiment than in the second year (Fig 4). This can be explained by the fact that actual plant density was generally lower in the first vs. second year (Fig 2) and by the negative correlation between actual plant density and PC (Fig 6b). Still, PC was relatively higher than in quinoa grains (Pereira et al., 2019): in a previous study conducted in Israel under similar conditions, quinoa grain PC ranged between 5.3% and 14.2% (Asher et al., 2022); in another study conducted in Vietnam, quinoa grain PC ranged between 15.01% and 18.49% [47]. In addition, the PC of young vegetative quinoa was relatively higher than in other vegetative crops, such as spinach, with a reported PC of between 19.9% and 22.8% [38]. Hence, when considering these data and the relatively high DM yield described above, young green quinoa may be cultivated in Israel and other Mediterranean countries as a potential source of plant-based protein. Supporting this notion is the amino acid composition of the young green quinoa protein (Tables 2 and 3). In most cases, the EAA composition in 100 g DM of young green quinoa fulfilled over 50% of the recommended daily intake of EAA for a 70 kg human adult [41]. In some cases, the EAA composition was above 100% of the recommended daily intake. These data suggest the high quality of young green quinoa protein because the nutritional quality of a protein is determined by its concentration of EAA, which humans cannot synthesize [32]. Considering that the protein requirement for human adults stands at 0.66 g kg^-1^ per day [41], 200 g DM of young green quinoa can meet the protein and most of the EAA requirements for a 70 kg adult. The volume of 100 g of dry young vegetative quinoa is 103 ml and the fresh matter weight is circa 900 g. This is of clear economic and social significance because the consumption of sufficient high-quality protein in the diet poses a challenge for some populations, particularly those who are limiting, or have eliminated their animal protein intake and must rely on cereals, grains, and legumes for their protein consumption [13].

Plant density is a critical factor affecting quinoa and other commercial field crops’ yield and quality traits [48]. In this study, it was interesting to find that plant density differed in the plots (Fig 2), possibly because of variation in seed quality, inaccuracies in the sowing operation or failed seedling development due to biotic or abiotic stress conditions [39,49]. However, these assumptions cannot be established in this present study, as germination studies were not conducted. Young vegetative quinoa DM and protein yields were both positively correlated to plant density at harvest (Fig 6a, c). In contrast, a study conducted in India showed that quinoa foliage yield generally increases with increasing row spacing [42]. Thus, the optimal plant density should be carefully considered because low plant density may result in thicker quinoa stems [20], reducing their palatability for humans. However, high plant density may reduce plant quality, as in this study, young vegetative quinoa PC was negatively correlated to plant density at harvest (Fig 6b). Future research should address this issue. Additional future aspects should address the effect of plant density and sowing architecture on industrialization and practical field cultural operations of the crop.

In both years of the experiment, young green quinoa protein yield was greatest on the December sowing date (Fig 5). In addition to this advantage, December is better for sowing crops because weed control is simpler than in November. As the weeds have already germinated in December, they can be effectively eliminated by soil cultivation or a suitable non-residual herbicide. With the help of this weed-control strategy, new quinoa shoots can grow with little or no requirement for herbicides [23]. Thus, the results of this present study suggest that the December sowing date is more appropriate for cultivation in the experimental setting. Additional in-depth research is necessary to further investigate the effect of climate and seasonal parameters on young vegetative quinoa production and quality. This is important, particularly in the Mediterranean region where precipitation is expected to decrease and temperature to warm significantly beyond the global average [50]. Moreover, protein yield was among the highest in most cases, albeit not significantly so, for the accession Peppermint (Fig 5). Therefore, this accession seems more suitable for cultivation in the experimental location when considering young vegetative quinoa for protein yield. Additional studies are required to further investigate how different genotypes impact the quality and production of young vegetative quinoa in locations worldwide. Also, enhanced leaf size as well as plant biomass and protein accumulation in the vegetative growth stages could be desirable traits for quinoa breeding programs.

In previous studies conducted in Israel, quinoa grain yield reached 4,266 kg DM ha^-1^, whereas PC stood at 14.2% [20]. From these data, protein yield reached 605 kg ha^-1^. In the present study, conducted under similar conditions, the protein yield of young vegetative quinoa was about 50% lower (Fig 5). However, a significant advantage of cultivating young leafy quinoa is the short growing cycle from sowing to harvest, usually 60–80 days during the winter. In comparison, winter quinoa cultivation for grain production is much longer, 150–170 days. Therefore, multiple harvests of young vegetative quinoa or establishing a new cultivation on the same plot following harvest may produce a similar or greater protein yield than quinoa grown for grains. In addition, a shorter growing cycle means that the plants may avoid various stresses, such as extreme weather events [51,52] and the young vegetative quinoa can be cultivated between winter and summer yearly crop cycles, thus maximizing land and other agricultural resource usages. Finally, the young vegetative quinoa requires scant irrigation (< 100 m^3^ ha^-1^), making it highly suitable as a protein-rich crop for semi-arid and possibly even arid regions, where irrigation infrastructure is a resource that is in short supply [53].

## 5. Conclusions

In conclusion, it seems that according to our research hypothesis, there are high prospects for cultivating young vegetative quinoa in Israel and other Mediterranean countries as a new sustainable protein-rich winter leafy crop. The fact that quinoa had high protein yield and quality, even though the plants received only a small amount of irrigation (100 m^3^ ha^-1^), supports this notion. This is crucial for agricultural production in arid and semi-arid regions where access to irrigation water is challenging and expensive. Nevertheless, further studies are required to examine other agrotechnical parameters, such as row spacing, plant density, irrigation and fertilization regimes, and their effects on yield and quality parameters. Studies should also evaluate the possibility of several growing cycles of young vegetative quinoa on the same plots with different sowing dates to maximize land use and yearly yield potential. Also, winter temperatures in the Mediterranean climate may drop below 0⁰ C (Table 1). Although quinoa is well known for its high level of frost resistance, convective frosts may result in necrotic spots in the foliage [54]. Thus, the impact of freezing temperatures and frost events should be assessed and considered when cultivating young vegetative quinoa during cold winters. Finally, evaluating its potential as a short-cycle high-quality forage crop and various economic aspects such as production cost will enable young vegetative quinoa’s geographical distribution and expansion.

## Acknowledgements

The authors thank Shaul Graph, Itzik Abarbanel and the Avnei-Eitan research farm team for their efforts in performing this study. The ICA foundation in Israel funded several aspects of this study.

## Supporting information

S1 file. Data tables for figures 2-5. (XSLX)

## References

1. Grandview research. Global Protein Ingredients Market Size Report, 2021-2028. 2020 [cited 29 Jan 2023]. Available: https://www.grandviewresearch.com/industry-analysis/protein-ingredients-market

2. de Boer J, Aiking H. On the merits of plant-based proteins for global food security: Marrying macro and micro perspectives. Ecological Economics. 2011. pp. 1259–1265. doi:10.1016/j.ecolecon.2011.03.001

3. Day L, Cakebread JA, Loveday SM. Food proteins from animals and plants: Differences in the nutritional and functional properties. Trends in Food Science and Technology. Elsevier Ltd; 2022. pp. 428–442. doi:10.1016/j.tifs.2021.12.020

4. Pam Ismail B, Senaratne-Lenagala L, Stube A, Brackenridge A. Protein demand: Review of plant and animal proteins used in alternative protein product development and production. Animal Frontiers. 2020;10: 53–63. doi:10.1093/af/vfaa040

5. Aschemann-Witzel J, Gantriis RF, Fraga P, Perez-Cueto FJA. Plant-based food and protein trend from a business perspective: markets, consumers, and the challenges and opportunities in the future. Critical Reviews in Food Science and Nutrition. Taylor and Francis Inc.; 2020. pp. 1–10. doi:10.1080/10408398.2020.1793730

6. Andreotti F, Bazile D, Biaggi C, Callo-concha D, Jacquet J, Jemal OM, et al. When neglected species gain global interest: Lessons learned from quinoa’s boom and bust for teff and minor millet. Glob Food Sec. 2022;32: 100613. doi:10.1016/j.gfs.2022.100613

7. Bazile D, Jacobsen S-E, Verniau A. The Global Expansion of Quinoa: Trends and Limits. Front Plant Sci. 2016;7: 1–6. doi:10.3389/fpls.2016.00622

8. Bazile D. (ed.), Bertero H.D. (ed.), Nieto C. (ed.). State of the art report on quinoa around the world in 2013. Santiago de Chili: FAO; CIRAD; 2015. pp. 603. https://www.fao.org/quinoa-2013/publications/detail/en/item/278923/icode/?no_mobile=1

9. Choukr-Allah R, Rao NK, Hirich A, Shahid M, Alshankiti A, Toderich K, et al. Quinoa for marginal environments: Toward future food and nutritional security in MENA and central Asia regions. Frontiers in Plant Science. Frontiers Research Foundation; 2016. doi:10.3389/fpls.2016.00346

10. Pulvento C, Bazile D. Worldwide Evaluations of Quinoa—Biodiversity and Food Security under Climate Change Pressures: Advances and Perspectives. Plants. 2023;12: 868. doi:10.3390/plants12040868

11. Jacobsen SE. The scope for adaptation of quinoa in Northern Latitudes of Europe. J Agron Crop Sci. 2017;203: 603–613. doi:10.1111/jac.12228

12. Alandia G, Rodriguez JP, Jacobsen SE, Bazile D, Condori B. Global expansion of quinoa and challenges for the Andean region. Glob Food Sec. 2020;26. doi:10.1016/j.gfs.2020.100429

13. Scanlin L, Lewis KA. Quinoa as a Sustainable Protein Source: Production, Nutrition, and Processing. Sustainable Protein Sources. Elsevier Inc.; 2017. pp. 223–238. doi:10.1016/B978-0-12-802778-3.00014-7

14. Pereira E, Encina-Zelada C, Barros L, Gonzales-Barron U, Cadavez V, C.F.R. Ferreira I. Chemical and nutritional characterization of *Chenopodium quinoa* Willd (quinoa) grains: A good alternative to nutritious food. Food Chem. 2019;280: 110–114. doi:10.1016/J.FOODCHEM.2018.12.068

15. Angeli V, Silva PM, Massuela DC, Khan MW, Hamar A, Khajehei F, et al. Quinoa (*Chenopodium quinoa* Willd.): An overview of the potentials of the “golden grain” and socio-economic and environmental aspects of its cultivation and marketization. Foods. 2020;9. doi:10.3390/foods9020216

16. Ceyhun Sezgin A, Sanlier N. A new generation plant for the conventional cuisine: Quinoa (*Chenopodium quinoa* Willd.). Trends Food Sci Technol. 2019;86: 51–58. doi:10.1016/J.TIFS.2019.02.039

17. Dakhili S, Abdolalizadeh L, Hosseini SM, Shojaee-Aliabadi S, Mirmoghtadaie L. Quinoa protein: Composition, structure and functional properties. Food Chemistry. Elsevier Ltd; 2019. doi:10.1016/j.foodchem.2019.125161

18. Noulas C, Tziouvalekas M, Vlachostergios D, Baxevanos D, Karyotis T, Iliadis C. Adaptation, Agronomic Potential, and Current Perspectives of Quinoa Under Mediterranean Conditions: Case Studies from the Lowlands of Central Greece. Commun Soil Sci Plant Anal. 2017;48: 2612–2629. doi:10.1080/00103624.2017.1416129

19. Sellami MH, Pulvento C, Lavini A. Agronomic practices and performances of quinoa under field conditions: A systematic review. Plants. 2021;10: 1–20. doi:10.3390/plants10010072

20. Asher A, Dagan R, Galili S, Rubinovich L. Effect of Row Spacing on Quinoa (Chenopodium quinoa) Growth, Yield, and Grain Quality under a Mediterranean Climate. Agriculture. 2022;12: 1298. doi:10.3390/agriculture12091298

21. Walters H, Carpenter-Boggs L, Desta K, Yan L, Matanguihan J, Murphy K. Effect of irrigation, intercrop, and cultivar on agronomic and nutritional characteristics of quinoa. Agroecology and Sustainable Food Systems. 2016;40: 783–803. doi:10.1080/21683565.2016.1177805

22. Wang N, Wang F, Shock CC, Meng C, Qiao L. Effects of management practices on quinoa growth, seed yield, and quality. Agronomy. 2020;10. doi:10.3390/agronomy10030445

23. Asher A, Galili S, Whitney T, Rubinovich L. The potential of quinoa (*Chenopodium quinoa*) cultivation in Israel as a dual-purpose crop for grain production and livestock feed. Sci Hortic. 2020;272: 109534. doi:10.1016/j.scienta.2020.109534

24. Filik G. Biodegradability of quinoa stalks: The potential of quinoa stalks as a forage source or as biomass for energy production. Fuel. 2020;266: 117064. doi:10.1016/j.fuel.2020.117064

25. Matías J, Cruz V, Reguera M. Heat Stress Impact on Yield and Composition of Quinoa Straw under Mediterranean Field Conditions. Plants. 2021;10: 955. doi:10.3390/PLANTS10050955

26. Ebeid HM, Kholif AE, El-Bordeny N, Chrenkova M, Mlynekova Z, Hansen HH. Nutritive value of quinoa (*Chenopodium quinoa*) as a feed for ruminants: in sacco degradability and in vitro gas production. Environmental Science and Pollution Research. 2022;29: 35241–35252. doi:10.1007/s11356-022-18698-x

27. Ramos N, Cruz AM. Evaluation of seven seasonal crops for forage production during the dry season in Cuba. Cuban Journal of Agricultural Science. 2002;36: 271–276.

28. Camaggio G, Amicarelli V. The ancient crop of quinoa for world food security. In: Mis’niakiewicz M, Popek S, editors. Future Trends and Challenges in the Food Sector. Cracow, Poland: Polish Society of Commodity Science; 2014.

29. Pathan S, Siddiqui RA. Nutritional Composition and Bioactive Components in Quinoa (*Chenopodium quinoa* Willd.) Greens: A Review. Nutrients. 2022;14: 1–12. doi:10.3390/nu14030558

30. Adamczewska-Sowińska K, Sowiński J, Jama-Rodzeńska A. The effect of sowing date and harvest time on leafy greens of quinoa (*Chenopodium quinoa*) willd.) yield and selected nutritional parameters. Agriculture (Switzerland). 2021;11. doi:10.3390/agriculture11050405

31. Vazquez-Luna A, Cortés VP, Carmona FF, Díaz-Sobac R. Quinoa leaf as a nutritional alternative. Cienc Investig Agrar. 2019;46: 137–143. doi:10.7764/rcia.v46i2.2098

32. Pathan S, Eivazi F, Valliyodan B, Paul K, Ndunguru G, Clark K. Nutritional Composition of the Green Leaves of Quinoa (*Chenopodium quinoa* Willd.). J Food Res. 2019;8: 55. doi:10.5539/jfr.v8n6p55

33. Gawlik-Dziki U, Swieca M, Sulkowski M, Dziki D, Baraniak B, Czyz J. Antioxidant and anticancer activities of *Chenopodium quinoa* leaves extracts - In vitro study. Food and Chemical Toxicology. 2013;57: 154–160. doi:10.1016/j.fct.2013.03.023

34. Stoleru V, Jacobsen SE, Vitanescu M, Jitareanu G, Butnariu M, Munteanu N, et al. Nutritional and antinutritional compounds in leaves of quinoa. Food Biosci. 2022;45. doi:10.1016/j.fbio.2021.101494

35. Lim JG, Park HM, Yoon KS. Analysis of saponin composition and comparison of the antioxidant activity of various parts of the quinoa plant (*Chenopodium quinoa* Willd.). Food Sci Nutr. 2020;8: 694–702. doi:10.1002/fsn3.1358

36. Dick Mastebroek H, Limburg H, Gilles T, Marvin HJ. Occurrence of sapogenins in leaves and seeds of quinoa (Chenopodium quinoa Willd).

37. Bernardo Solíz-Guerrero J, Jasso De Rodriguez D, Rodríguez-García R, Luis Angulo-Sánchez J, Méndez-Padilla G. Quinoa Saponins: Concentration and Composition Analysis. ASHS Press; 2002.

38. Abd El-Samad EH, Hussin SA, El-Naggar AM, El-Bordeny NE, Eisa SS. The potential use of quinoa as a new non-traditional leafy vegetable crop. Bioscience Research. 2018;15: 3387–3403. Available: www.isisn.org

39. Bellalou A, Daklo-Keren M, Abu Aklin W, Sokolskaya R, Rubinovich L, Asher A, et al. Germination of *Chenopodium quinoa* cv. ‘Mint Vanilla’ seeds under different abiotic stress conditions. Seed Science and Technology. 2022;50: 41–45. doi:10.15258/sst.2022.50.1.05

40. El-Naggar A, Hussin S, Abd El-Samad E, Eisa S. Quinoa As a New Leafy Vegetable Crop in Egypt. Arab Universities Journal of Agricultural Sciences. 2018;26: 745–753. doi:10.21608/ajs.2018.16007

41. WHO. Protein and amino acid requirements in human nutrition: report of a joint WHO/FAO/UNU expert consultation. 2007.

42. Bhargava A, Sudhir S, Deepak O. Effect of sowing dates and row spacings on yield and quality components of quinoa (*Chenopodium quinoa*) leaves. Indian Journal of Agricultural Sciences. 2007;77: 748–751. Available: https://www.researchgate.net/publication/215713006

43. Woodard KR, Prine GM. Forage Yield and Nutritive Value of Elephantgrass as Affected by Harvest Frequency and Genotype (AJ). Agron J. 1991;83: 541–546.

44. Robins JG, Bauchan GR, Brummer EC. Genetic mapping forage yield, plant height, and regrowth at multiple harvests in tetraploid Alfalfa (*Medicago sativa* L.). Crop Sci. 2007;47: 11–18. doi:10.2135/cropsci2006.07.0447

45. Bantis F, Kaponas C, Charalambous C, Koukounaras A. Strategic successive harvesting of rocket and spinach baby leaves enhanced their quality and production efficiency. Agriculture (Switzerland). 2021;11. doi:10.3390/agriculture11050465

46. Benlhabib O, Yazar A, Qadir M, Lourenço E, Jacobsen SE. How can we improve Mediterranean cropping systems? J Agron Crop Sci. 2014;200: 325–332. doi:10.1111/jac.12066

47. van Minh N, Hoang DT, van Loc N, Long NV. Effects of plant density on growth, yield and seed quality of quinoa genotypes under rain-fed conditions on red basalt soil regions. Aust J Crop Sci. 2020;14: 1977–1982. doi:10.21475/ajcs.20.14.12.2849

48. Spehar CR, Rocha JE da S. Effect of sowing density on plant growth and development of quinoa, genotype 4.5, in the Brazilian Savannah Highlands. Bioscience Journal. 2009;25: 53–58.

49. Jacobsen S-E. The Worldwide Potential for Quinoa (*Chenopodium quinoa* Willd.). Food Reviews International. 2003;19: 167–177. doi:10.1081/FRI-120018883

50. Lionello P, Scarascia L. The relation between climate change in the Mediterranean region and global warming. Reg Environ Change. 2018;18: 1481–1493. doi:10.1007/s10113-018-1290-1

51. Lakshmanan P, Robinson N. Stress Physiology: Abiotic Stresses. Sugarcane: Physiology, Biochemistry, and Functional Biology. 2013; 411–434. doi:10.1002/9781118771280.CH16

52. Wahbi A, Sinclair TR. Simulation analysis of relative yield advantage of barley and wheat in an eastern Mediterranean climate. Field Crops Res. 2005;91: 287–296. doi:10.1016/j.fcr.2004.07.020

53. Fuentes F, Bhargava A. Morphological Analysis of Quinoa Germplasm Grown Under Lowland Desert Conditions. J Agron Crop Sci. 2011;197: 124–134. doi:10.1111/j.1439-037X.2010.00445.x

54. Jacobsen SE, Monteros C, Corcuera LJ, Bravo LA, Christiansen JL, Mujica A. Frost resistance mechanisms in quinoa (*Chenopodium quinoa* Willd.). European Journal of Agronomy. 2007;26: 471–475. doi:10.1016/j.eja.2007.01.006

